# Improved genomic resources for the swordtail cricket, *Laupala kohalensis* Otte 1994

**DOI:** 10.64898/2026.06.02.729584

**Authors:** Nicholai M. Hensley, Benjamin A. Sandkam, Kerry L. Shaw

**Affiliations:** Department of Neurobiology and Behavior, Cornell University, Ithaca, New York, USA; Department of Zoology, University of Cambridge, Cambridge, Cambridgeshire, United Kingdom

**Author notes:** Corresponding author: Nicholai M. Hensley.

**Keywords:** non-adaptive radiation, speciation, sexual selection, Orthoptera, Hawaii, adaptive radiation

## Abstract

Advances in genetic tools such as next and third generation sequencing, paired with a focus on representative clades, provide insight into how processes including adaptation, admixture, and genome structure shape the evolution and maintenance of species. However, our understanding of the genomics of speciation is dominated by systems where ecological adaptations are thought to cause initial barriers to gene exchange. In contrast to other model systems, the 38 species of the genus *Laupala* constitute a very rapid radiation, where evolution of reproductive barriers and speciation is thought to be driven by sexual selection. Here, with novel PacBio HiFi reads and RNA- and Iso-Seq data, we provide a highly contiguous, chromosome-level genome and markedly improved annotation of the endemic Hawaiian cricket*, Laupala kohalensis* Otte, 1994. Our new resources advance previous efforts, placing 99% of 47 scaffolds on 7 autosomes and 1 sex chromosome in the 1.67 Gb assembly, with a 98.8% BUSCO score (insecta_db10), N50 of ∼268 Mb, and L50 of 3. Using a custom repeat library, we estimate the genome to have 46.09% repeat content, and the new annotation includes an increased estimate of 17,670 genes, which coincides with that known from other Orthopterans. Notably, we find a large nuclear DNA segment of mitochondrial origin on chromosome 7. This new resource provides a powerful tool to identify and compare genomic causes of phenotypic diversification in a system characterized by strong signatures of sexual differentiation, representing an underappreciated but potentially widespread cause of speciation.

## Introduction

A central goal in evolutionary biology is uncovering the genetic mechanisms fueling the diversity of life. Next generation sequencing has facilitated data collected towards this goal. These efforts have revealed insights into how deterministic and parallel the mechanisms of speciation can be (Soria-Carrasco et al. 2014), and even how reticulate this process is (Meier et al. 2023). Because most species diversity accumulates as a consequence of species radiations (Wiens and Moen 2025), much attention has focused on clades with changes in diversification rate (Scholl and Wiens 2016). Moreover, many of these focal clades possess well-described ecological adaptations (Schluter 2000), suggesting the role of natural selection on ecological trait differentiation in speciation. However, a balanced examination of speciation processes should also include ecologically conserved groups, to reveal general principles concerning the mechanisms that drive variation in diversity (Rundell and Price 2009).

The genus *Laupala* Otte 1994 (Otte 1994) has emerged as a model clade for speciation research, characterized by a rapid speciation rate largely attributed to non-ecological selection. Other members of the swordtail cricket family (Trigonidiidae) are diverse and geographically widely distributed, with (generally) small-bodied species whose females possess sabre-like ovipositors. *Laupala* species are single-island endemics, found only on the volcanic islands of Hawaii and have radiated rapidly across the archipelago, resulting in 38 known species (Otte 1994; Shaw 2000). Like many crickets and their allies, *Laupala* males sing by stridulating their forewings at characteristic rates to produce a relatively simple song composed of single, pure-tone notes ∼5000 Hz (Fig. 1A). Females respond to male calling song via phonotaxis, and then begin a complex, prolonged courtship including antennation (facilitating olfactory or gustatory signaling (Mullen et al. 2007)), reciprocal facing, the transfer of multiple, spermless, microspermatophores, and ending with the transfer of a sperm-filled macrospermatophore (deCarvalho and Shaw 2005; Shaw and Khine 2004). Female preferences exert directional selection on male songs (Oh and Shaw 2013), male and female acoustic behaviors are phenotypically and genetically correlated (Grace and Shaw 2011, 2012; Xu and Shaw 2026), and have diverged within and between species (Mendelson and Shaw 2005; Blankers et al. 2018, 2019), suggesting that sexual selection on pre-mating behaviors like song is strong in this system.

**Figure 1.**
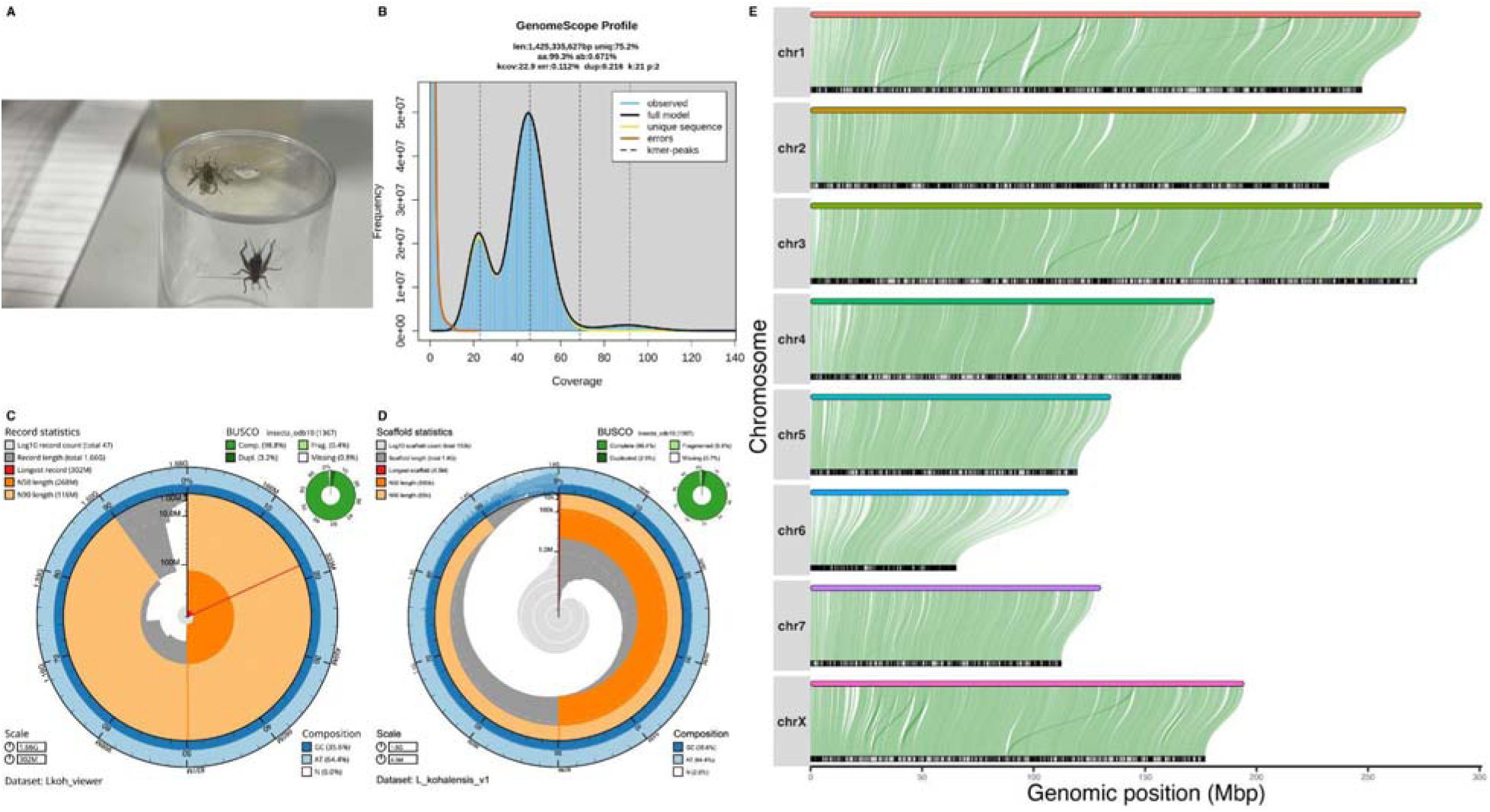
(A) Image of a male *Laupala kohalensis* stridulating with raised wings to a female; note the curved ovipositor between the cerci at the base of her abdomen. (B) GenomeScope profile of the raw PacBio HiFi reads. (C,D) Snail plots comparing the (C) current and (D) previous genome assemblies for *L. kohalensis*. (E) Contigs of the previous genome mapped (lower tracks) to the largest scaffolds (chromosomes) of the new genome (upper tracks), showing high contiguity between the two, and high completeness of the current assembly. This is an improvement from the previous assembly, which had ∼50% unplaced scaffolds (See Fig. S1). Contigs limited to those 10,000 bp or longer for visualization.

To better explore the genetic basis of phenotypic differentiation as it relates to barriers to gene exchange in this group, and speciation more broadly, we developed a chromosome-level genome for the cricket *Laupala kohalensis* from the Big Island of Hawaii, improving upon a previous version (Blankers et al. 2018), and present a new genome annotation using *Laupala* - specific RNA- and Iso-seq data. Males of *L. kohalensis* have one of the fastest songs known among *Laupala* species, and quantitative trait locus mapping with hybrids between *L. kohalensis* and the slower singing *L. paranigra* has localized differences in male song to 8-10 loci of relatively small effect sizes (Shaw et al. 2007; Waller et al. 2023). Many of these regions co-localize with female preferences for differences in male song, indicating that tight genetic linkage or pleiotropy connect the changes in acoustic signals with their respective preferences (Wiley et al. 2012; Xu and Shaw 2019, 2026, 2021; Shaw and Lesnick 2009). Although much progress has been made to link genes and behavior to the rapid speciation in this group, the previous genome was highly fragmented with ∼50% of scaffolds lacking placement to any linkage group. Furthermore, the only available annotation was produced largely by consulting gene evidence from a distantly related genus (Ylla et al. 2021).

Made with PacBio long read data, our *de novo* genome improves the contiguity of the previous assembly and doubles the amount of sequence placed within linkage groups, creating a chromosome-level (n = 8 chromosomes) assembly. The assembly size is 1.67 Gb, with 90% contained on 8 scaffolds and possesses an estimated 46% repetitive content. As with other Orthopteran species, we also show evidence for a near complete mtDNA genome, along with nuclear mitochondrial DNA segments (NUMTs). We generate a new annotation containing 17,670 genes. With better gene models and higher contiguity, the revised genome of *Laupala kohalensis* will enable studies on speciation genomics within the clade and comparative genomics with other Arthropod groups.

## Methods

### Biological materials

Individual *L. kohalensis* were collected as juveniles and adults in 2003 from the Kohala Mountains, Hawaii, USA (North Kohala District, Hawaii; elevation 370-380 m; latitude 20.182848, longitude -155.729173). Full sib-matings from the progeny of a single female were used to generate a stock colony, which has subsequently undergone many generations (25+) of inbreeding and is maintained in the Shaw lab at Cornell University.

### DNA extraction for PacBio HiFi library preparation & Revio sequencing

A single, live male was frozen whole, overnight at -80*C for DNA isolation the following day. To meet tissue amount requirements, we separated the head and thorax from the abdomen using a clean pipette tip and placed each half into separate 1.5 mL Eppendorf tubes. We used a microbalance to weigh each half to determine the maximum amount of tissue to use for extraction (14.4 ug). We extracted HMW DNA from the abdomen using the NEB Monarch Extraction Kit for Tissue, with slight modifications for yield improvement. Notably, we used a high salt procedure in order to precipitate out strands of HMW DNA. After quantifying yield on a Nanodrop, DNA was sent to Cornell Weill School of Medicine Genome center for sequencing.

### DNA sequencing, genome assembly, and scaffolding

We sequenced the sample on a PacBio Revio machine using one flow cell to generate 4,919,653 PacBio HiFi reads. Total run time was 46.5 hours, producing 144 Gb. Assembly proceeded with five separate rounds of hifiasm (Cheng et al. 2021) to test assembly sensitivity to varying parameters, including defaults. From the most contiguous assembly, we then used BlobTools (Laetsch and Blaxter 2017) to screen for any contigs that were the result of potential contamination and purge them from the assembly. We used Inspector (Chen et al. 2021) to compare the raw reads to the preliminary contig assembly to diagnose and fix small structural variants. We then used markers from a previously published linkage mapping study (Blankers et al. 2018) to scaffold the contigs, placing 500 N’s between contig joins. Scaffolds were assigned chromosome names according to the linkage map grouping, and in line with the previous assembly. One contig mapped to multiple scaffolds (corresponding to two chromosomes) and manual inspection of the read suggested a misassembly of that single contig, which we subsequently split before re-scaffolding. We used GenomeScope (Vurture et al. 2017) to check the distribution of k-mers on the raw reads, BUSCO (Manni et al. 2021) to assess genome quality at various stages in the assembly pipeline (to compare different assemblies during the purging of duplicates, pre-scaffolding, and post-scaffolding), and QUAST (Gurevich et al. 2013) during the initial and final stages of assembly. For list of all computational tools used our analyses, see Table 1.

### Identification of mitochondrial DNA sequence

To identify mitochondrial DNA in our new assembly, we used MitoHiFi (Uliano-Silva et al. 2023). Preliminary attempts using the final assembly indicated that multiple, large regions of high similarity might be present throughout the assembly. Such patterns are expected with nuclear copies of the mitochondrial DNA (NUMTs), which have been identified in other Orthoptera genomes (Liu, Liu, et al. 2024; Song et al. 2014). Instead, we used MitoHiFi on the raw reads to identify the entire mitochondrial sequence (mtDNA). We then added this sequence back to the final genome assembly. We aligned the newly identified mtDNA sequence to the assembly using minimap2 (Li 2018, 2021) to locate other regions of the genome assembly with high similarity, indicative of a NUMT. To confirm the presence of a NUMT instead of assembly error, we mapped all the trimmed reads to both the final assembly with the MitoHiFi mtDNA and the final assembly without the identified MitoHifi mtDNA added back to look at the change in read depth across the NUMT region. If NUMTs are present in the assembly without the MitoHiFi identified mtDNA contig, we expect such regions to have high mapping rates with the trimmed read data as true mtDNA read copies should be high and able to map to other regions with high sequence similarity. However, in an assembly with the mtDNA properly identified and added back to the genome, we expect that any mtDNA reads previously mapping incorrectly to the NUMT will be subsequently mapped to the actual mtDNA contig now present in the assembly. To map reads in both versions and compare areas where read-depth changed due to the presence of the mtDNA-like sequence within the assembly, we used mosdepth (Pedersen and Quinlan 2018) to count read depth across regions in 1000 bp sliding windows created by bedtools (Quinlan 2014) and that overlap by 500 bp.

### RNA extraction for short-read RNA-seq library preparation and NovaSeq 6000 sequencing

Samples of 8 males were homogenized with RNAse free sterile pestles inside individual 1.5 mL Eppendorf collection tubes. We extracted total RNA from full body samples with the RNeasy Mini Prep Kit (Qiagen). Quantification and quality control were performed on an Agilent 2100 BioAnalyzer in the Cornell University Biotechnology Core Facility. Each sample was then individually indexed and prepared for sequencing after Poly(A) selection and sequenced on an Illumina NovaSeq 6000 platform as 150 bp PE reads by Azenta Life Sciences (South Plainfield, NJ, USA).

### RNA extraction for ISO-seq library preparation and NovaSeq 6000 sequencing

To improve gene models during annotation, we generated PacBio HiFi reads of full-length RNA isoforms (Iso-seq). We pooled whole body tissues from two adult males, two adult females, one juvenile male, and one juvenile female to maximize isoform diversity before extracting RNA using a Monarch Total RNA Miniprep Kit (New England Biolabs). RNA quality and quantity were verified using a NanoDrop One and Qubit (ThermoScientific), and RNA integrity was verified on a Fragment Analyzer (RQN = 8.1).

The reads were generated in multiplex alongside seven samples from an unrelated project by incorporating unique molecular identifiers (UMI) during library preparation with the Kinnex full-length RNAseq library prep kit and sequenced on PacBio Revio (Pacific BioSciences). RNA libraries were split, demultiplexed, and primers/poly-A tails removed using lima (v 2.9.0) (Lima Docs, n.d.) and skera (v 1.2.0) (Skera Docs, n.d.).

### Repeat masking & genome annotation

We used a combination of approaches to annotate the genome. First, we generated a custom repeat library using RepeatModeler (Flynn et al. 2020) and RepeatMasker following (Bowman, n.d.). Briefly, to use in RepeatModeler, we built a custom repeat database with the following resources and tools, which were then combined and used to scan the genome for similar elements: (1) the Repbase (Bao et al. 2015) repeat element libraries for *Drosophila*, *Anopheles*, Invertebrates, and Invertebrate Supplement; (2) the library of Insecta short-interspersed nuclear elements (SINEs) from SINEBase (Vassetzky and Kramerov 2013); (3) the Dfam transposable element (TE) library for arthropods (libraries 1,4,7,8) (Storer et al. 2021); (4) scanning the assembly for transposons with Transposon-PSI (“TransposonPSI: An Application of PSI-Blast to Mine (Retro-)Transposon ORF Homologies,” n.d.); (5) scanning the assembly for miniature inverted TEs with MITE-Tracker (Crescente et al. 2018); (6) scanning the assembly for long terminal repeats with LTR-Harvest (Ellinghaus et al. 2008) and LTR-Digest (Steinbiss et al. 2009); and (7) an Orthopteran specific repeat library (Liu, Zhao, et al. 2024). We combined these databases and sequences identified by scans into a merged library, removed short and redundant sequences with seqtk (Li, n.d.), and classified them with RepeatClassifier. After classification, we identified via BLAST-X (Camacho et al. 2009) and subsequently removed sequences from the “Unknown” category that were similar to proteins in the Uniprot Insecta database; we removed these sequences from the repeat library to prevent their masking. The final custom repeat library contained 3,182 sequences that were used by RepeatMasker to soft-mask the genome for annotation.

For genome annotation we performed two parallel rounds using BRAKER3 (Gabriel et al. 2024) that differed in the RNA evidence, as a combination protocol using both short-read and long-read sequencing was not widely tested and available at the time. Regardless of which RNA data, we ran BRAKER3 with a combined library of protein sequences as evidence in both parallel runs. The previous version of the *L. kohalensis* genome was annotated with limited RNA evidence (Ylla et al. 2021), but the content of which may still be informative for gene modeling. We used LiftOff (Shumate and Salzberg 2019) to map coordinates between the new assembly and the genes within the annotation from the previous assembly. We fed these gene models to the TAGADA pipeline (Kurylo et al. 2023), which can update them using the new genome and RNA-seq data; we used both the previously published (Waller et al. 2023) and our newly generated short-read RNAseq data to update the gene models. From these updated models, we translated protein sequences with TransDecoder (*TransDecoder: TransDecoder Source*, n.d.) and combined these subsequent proteins with the Arthropod sequences in OrthoDB (University of Greifswald 2018) to use as protein evidence in both runs of BRAKER3 (Gabriel et al. 2024).

In one run, we used the aforementioned protein evidence and both the published and the new, short-read RNAseq as evidence for the gene modeling. Short-reads were trimmed using TRIM-Galore (Krueger, n.d.) before use in BRAKER3. From the RNAseq data we aligned 863,621,150 reads from 12 RNA-seq samples. This run produced 15,186 gene models. In a separate instance of BRAKER3, we used both the protein evidence and the Iso-seq data for gene modeling. From our IsoSeq data we aligned 314,435 high-quality full-length RNA reads. This run produced 16,031 gene models. Afterwards, the annotations were combined using TSEBRA (Gabriel et al. 2021) into a final annotation with 17,670 genes (Table 2).

After genome annotation, we added functional annotations to the gene models using InterProScan (Paysan-Lafosse et al. 2023). We used eggNOG (Cantalapiedra et al. 2021; Huerta-Cepas et al. 2019) to add GO terms. To add symbols to genes, we downloaded the *Drosophila melanogaster* proteome from UniProt (UniProt Consortium 2023) and used blast-p matches at 1e-6 or lower to provide gene names to models within the *L. kohalensis* annotation. We used AGAT (Dainat et al. 2026) to combine the results from these three functional annotation steps to produce the final annotation. We renamed contigs to their scaffold positions on chromosomes using a custom script, and used BUSCO to check the annotation completeness.

To annotate the mitochondrial genome, we uploaded the sequence identified by MitoHiFi to the European Galaxy webportal at https://usegalaxy.eu/ (Galaxy Community 2024; “Galaxy,” n.d.) using MITOS2 (Donath et al. 2019; Al Arab et al. 2017), as well as using MFannot (Lang et al. 2023). We manually compared the annotations from different tools. We then performed the above functional annotation steps separately for the mtDNA annotation output by MITOS2, and then merged the mtDNA annotations with the rest of the genome annotation. We used AGAT for a number of transformations on the mtDNA annotation, including to manage functional annotations, and to merge and sort the annotations in their entirety. We visualized the annotation after manually confirming gene coordinates using the EZmito2 (Cucini et al. 2021) web server.

## Results

### Nuclear genome sequencing and assembly

PacBio HiFi sequencing produced 4,919,653 reads after trimming (min. length = 98 bp, mean length = 14584.3 bp, max. length = 56,322 bp), yielding an average of 43-fold coverage based on a genome size of 1.425 Gb from GenomeScope. Error rates were low (0.112%), and heterozygosity was moderately low (0.671%), which is expected given the sequenced individual was derived from a lab population maintained by full-sib inbreeding. The k-mer coverage histogram showed a tri-modal distribution with major peaks at 21.5- and 43-fold, and a minor bump at 86-fold coverage (Fig. 1B).

Our initial five assemblies with varying hifiasm parameters were highly similar, with 304 - 316 contigs. After choosing the most contiguous and purging duplicates, our assembly was 110 contigs and 1.66 Gb (Table 2), which is in-line with genome size estimates from the previous assembly (Blankers et al. 2018). This is larger than the GenomeScope estimate, most likely due to the repetitive nature of the genome and potentially residual heterozygous sequence, and smaller than estimates from nuclei staining in the congeneric *L. cerasina* (Petrov et al. 2000), which is diverged from *L. kohalensis* ∼ 1-2 Mya (Mendelson and Shaw 2005). Initial assembly correction by Inspector led to correction of 70 structural errors, including 41 expansions, 22 collapses, 4 switches, and 3 inversions; Inspector also corrected 10,916 small scale errors, most of which (8,782) were base substitutions. Scaffolding using data previously produced with recombination mapping (Blankers et al. 2018) led to a final nuclear genome of 46 scaffolds at a chromosome-level assembly, with 90% of contigs placed onto 7 autosomes and 1 sex chromosome separated by runs of 500 Ns (Fig. 1C). Even though the contigs from the long-read data were generally longer than the scaffolds produced in the previous genome assembly (Fig. 1D), the two versions are highly contiguous (Fig. 1E), confirming the recombination maps previously used for scaffolding are generally robust. Our chromosome-level assembly relied on relatively few recombination markers from these maps and only performed 63 joins to place the 110 contigs onto chromosomes. While the use of recombination mapping data does a generally good job of ordering contigs, it should be noted that there may still be minor misassemblies in the final order/orientation of contigs.

### Mitochondrial genome & detection of nuclear mtDNA

MitoHiFi produced a contiguous, circularized, and annotated mitochondrial genome which was 16,500 bp long (Fig. 2A,B) and that we merged back into the final assembly and annotation to produce a final assembly with 47 scaffolds on 8 chromosomes and 1 circular organelle genome. Comparing translations of annotated mitochondrial proteins produced by MitoHiFi, MITOS2, and MFannot revealed that coding positions were similar but generally less accurate with MitoHiFi, although tRNA sequences were identical across annotation methods.

**Figure 2.**
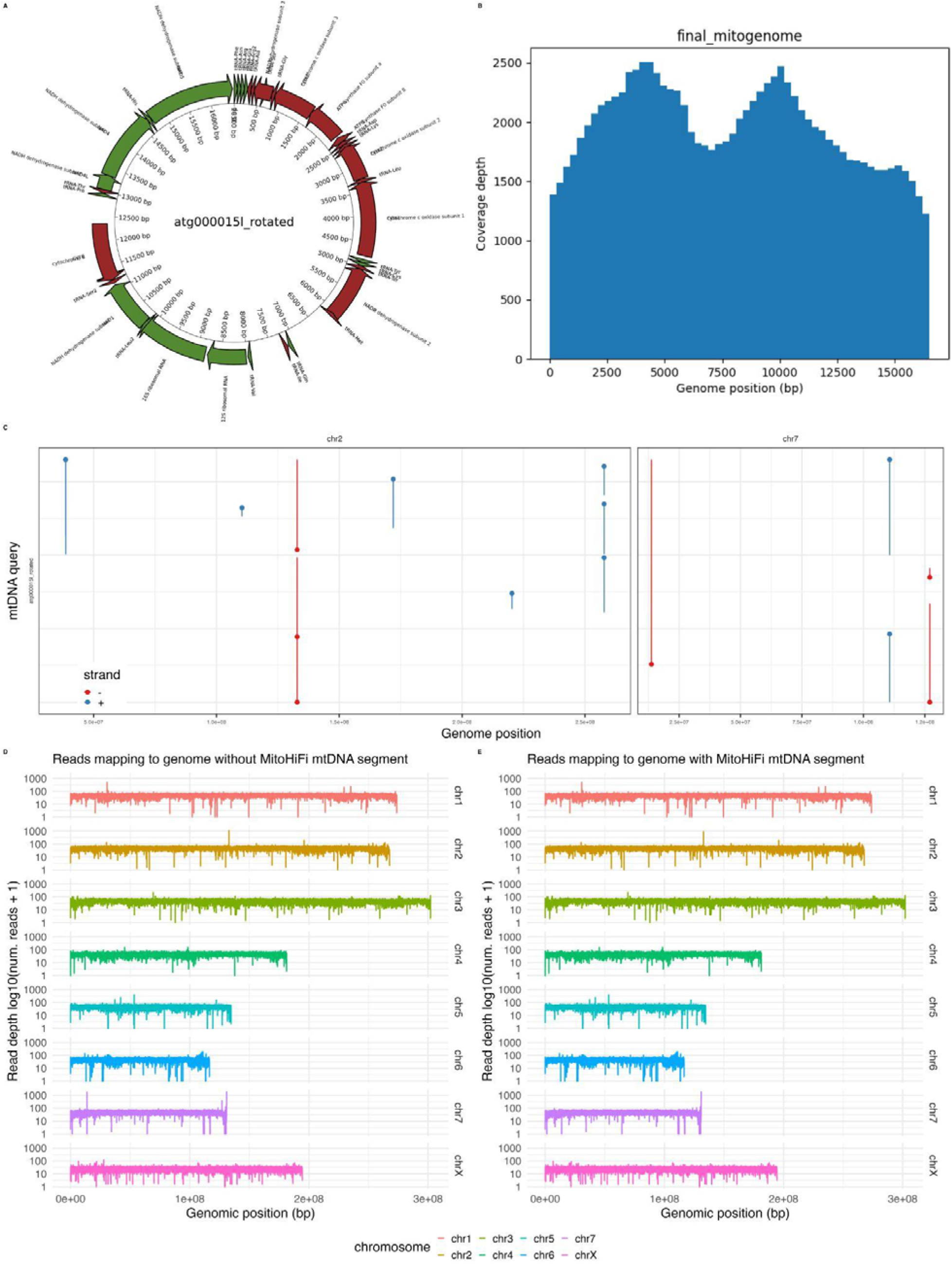
(A) Mitochondrial genome annotation from MFannot and (B) read coverage across the assembled mitogenome as provided by MitoHiFi. The mitochondrial sequence is included within the genome assembly as sequence atg000015l_rotated. (C) Nucmer alignment of the mtDNA sequence against the rest of genome assembly, highlighting multiple regions of similarity across two chromosomes (facets). Hits filtered to those 5500 bp or longer. Color is forward (red) or reverse strand (blue). (D,E) Read depth across all chromosomes in the final assembly without the mtDNA contig (D) and with the mtDNA contig (E). Note the two peaks on chromosome 7 with excessively high reads compared to chromosomal average; this first peak in read depth disappears and returns to baseline in the version of the assembly with the mtDNA contig (E). Y-axis is log10(x+1) transformed for visualization ease.

By aligning the mtDNA to the entire assembly with minimap2, we identified a large NUMT on chromosome 7 (Fig. 2C), as well as a number of smaller sequences with similarity to the identified mtDNA on other scaffolds. To ensure the NUMT was not an artifact of assembly, we looked at the number of reads crossing the NUMT join within the assembly, which was similar to the average genome coverage (43x) as expected if the region were truly part of the assembly. When we re-mapped the trimmed reads from the entire genome to both the assembly with the mtDNA (Fig. 2D) versus the assembly without the mtDNA sequence (Fig. 2E), we saw the read coverage dropped at that position to the average read coverage on chromosome 7, confirming that the mtDNA sequence pulled some but not all reads from that position on the assembly (Fig. 2E). Alignment of the NUMT to the mtDNA identified by MitoHiFi (Fig. S2) region indicated it is 13,921 bp long. Similar annotation of the NUMT region using MFannot showed that this sequence is generally intact compared to the identified mtDNA sequence, except it would theoretically lack genes for nad4 and tRNA-Histidine. As others have noted however, we do not expect NUMT sequences to produce functional transcripts, and out annotation served for comparison only (Hazkani-Covo et al. 2010).

### Identification and classification of repetitive elements

Using a custom repeat library and RepeatFinder, we found that 46.09% of the genome contains repeats, increased from an estimate of 35.51% in the original version of the genome. Such changes in repeat content are expected with longer reads (Wen et al. 2025; Peart et al. 2021; Liu et al. 2017; Cechova 2020); our new estimate is in agreement with other related orthopteran genomes (Szrajer et al. 2024; Kataoka et al. 2026; Ylla et al. 2021). RepeatFinder showed that 14.81% of repeats were retroelements, with 11.52% coming from long interspersed nuclear elements (LINEs), 0.00% coming from short interspersed nuclear elements (SINEs), and 3.29% coming from long terminal repeat elements (LTRs).

### Genome annotation

We soft masked 47.56% of the genome before annotation using the results from RepeatFinder in RepeatMasker. From AGAT summary statistics, the genome annotation produced 17,670 genes, 23,874 mRNA, 2 rRNA, and 23 tRNA models. The BUSCO score was 97.4%. See Table 2 for further details. Our annotation improves upon the number of genomic features from the previous annotation, which used less conspecific evidence and previously estimated 12,767 genes.

## Discussion

The *Laupala* genus is hyperdiverse, possessing 38 described species (Otte 1994; Shaw 2000) and the most rapid speciation rate known among invertebrates (Mendelson and Shaw 2005); this new genome and annotation will serve as the basis for understanding the genomics of speciation that underlies such a rapid radiation. Previously, (Blankers and Shaw 2024) used genotype-by-sequencing and the older *L. kohalensis* genome (Blankers et al. 2018) to understand the biogeographic and historical routes to divergence among populations of the Big Island endemic, *L. cerasina*. More recent work has leveraged a natural hybrid zone to identify patterns of genomic introgression between *L. makaio* and *L. orientalis (Sen and Shaw 2025)*. Both studies showcase how increased genomic data can shed light on the process of speciation and hybridization, broadly and within this system. A better resolved genome assembly and accompanying annotation will empower future work, including identifying genomic regions associated with barriers to gene flow and the genetic causes of phenotypic differentiation.

A scan of the current state of other Orthopteran genomes indicates that our resource is similar in quality to newer assemblies (Ylla et al. 2021; Szrajer et al. 2024; Kataoka et al. 2026). When surveying the larger families within Orthoptera (crickets, grasshoppers, and allies), genomic resources are unevenly distributed (Kataoka et al. 2022). As of April 2026, there were 42 reference-level Orthopteran genomes on NCBI; there are over 20,000 species of Orthoptera worldwide. Given the variation in genome size (Kataoka et al. 2022; Hawlitschek et al. 2023), diversity, habitat, and behavior (Song et al. 2020) within the order, this group represents an excellent system for evolutionary genomics if resources increase in taxonomic breadth.

We highlight evidence of a nuclear copy of the mitochondrial genome largely complete within *L. kohalensis*. Such NUMTs have previously been found in Orthopterans (Liu, Liu, et al. 2024; Song et al. 2014). Most studies relegate NUMTs to genomic fossils that hinder phylogenetic inference. Given their wide distribution across the Orthoptera, they may represent a unique opportunity to study the dynamics of genomic invasion. The functional role of these nuclear mitochondrial copies if any is generally unknown, although they have been implicated both in aging (Zhou et al. 2024) and disease (Hazkani-Covo et al. 2010; Wei et al. 2022) in other systems, as fragments of mitochondrial sequence can be inserted into and alter functional nuclear genes, themselves becoming functionally pseudogenized due to differences in codon usage between mitochondrial and nuclear genomes while effectively acting as indels.

Within *Laupala*, NUMT invasion may be both blessing and curse to understand speciation. Previous work before whole genome approaches show that the mtDNA within this clade does not possess the same branching pattern as the species relationships estimated from nuclear genomic material (Shaw 2002). However, these results may be sensitive to recent NUMT invasion if erroneously amplified sequences were included in the study. Our genome provides a resource to verify this result, and more broadly, quantify the variation in topology among different genomic regions during *Laupala* divergence. Conversely, depending on the length and similarity of NUMT to the mtDNA genome, they provide an orthogonal line of evidence to date the timing of divergence between lineages. The utility of NUMT regions in recreating phylogenetic relationships may be reduced if NUMTs are young and lineage-specific; including missing genes and gaps, our identified NUMT shares 84.17% similarity to the mtDNA along its aligned length (Fig. S2).

In producing a more contiguous genome with evidence-based annotations, we provide a resource for studying *Laupala* genomics as a counterpoint to systems with strong ecological adaptation driving speciation (Gillespie et al. 2020). As *Laupala* spp. are morphologically cryptic and ecologically indistinct (Hiller et al. 2019; Sen et al. 2025; Xu and Shaw 2020), they provide an excellent natural system to understand the genomic basis of species radiations writ large. Moreover, the contiguity of this improved genome and the new annotation can provide data for other comparative studies within Arthropoda (Song et al. 2020; Shin et al. 2024), such as exploring the stability or dynamism of genome organization across speciose Orthopterans (Zhang et al. 2025).

## Acknowledgements

Thanks to the staff at the Weill Cornell Medicine Genomics Core facility for their advice and help in preparing and sequencing a PacBio Revio library. The authors would also like to thank the staff at the Cornell BioHPC (Bukowski et al. 2010) for their generous time and support throughout this project.

## Data Availability

Data from this study are available under NCBI BioProject PRJNA1306088. PacBio HiFi raw sequencing for sample Lko2023_M1_from_Lko2003 are deposited in the NCBI Short Read Archive under SRR37501604. New RNA-seq reads are deposited at the SRA under SRR37605711, SRR37605710, SRR37605709, SRR37605708, SRR37605707, SRR37605706, SRR37605705, SRR37605704, and Iso-seq reads are deposited at the SRA as SRR37532061. The assembly CNL_Lkoh_2.0 has been deposited at GenBank under JBYTER000000000 (NCBI BioSample SAMN55867830). Scripts and notes for other analyses are at the GitHub repository: https://github.com/BSandkam/Laupala_kohalensis-Genome_Assembly_Hensley_et_al. The annotation and supplementary figures are also available on GitHub.

## Tables and Figures

Table 1. Software, version, any non-default parameter settings, and citation used for the assembly of a *de-novo Laupala kohalensis* genome, ordered by step within the pipeline.

Table 2. Statistics of the final genome assembly, the new genome annotation, and accessions for the sequence data and bioprojects produced during this project.

## Supplementary Material

**Figure S1.**
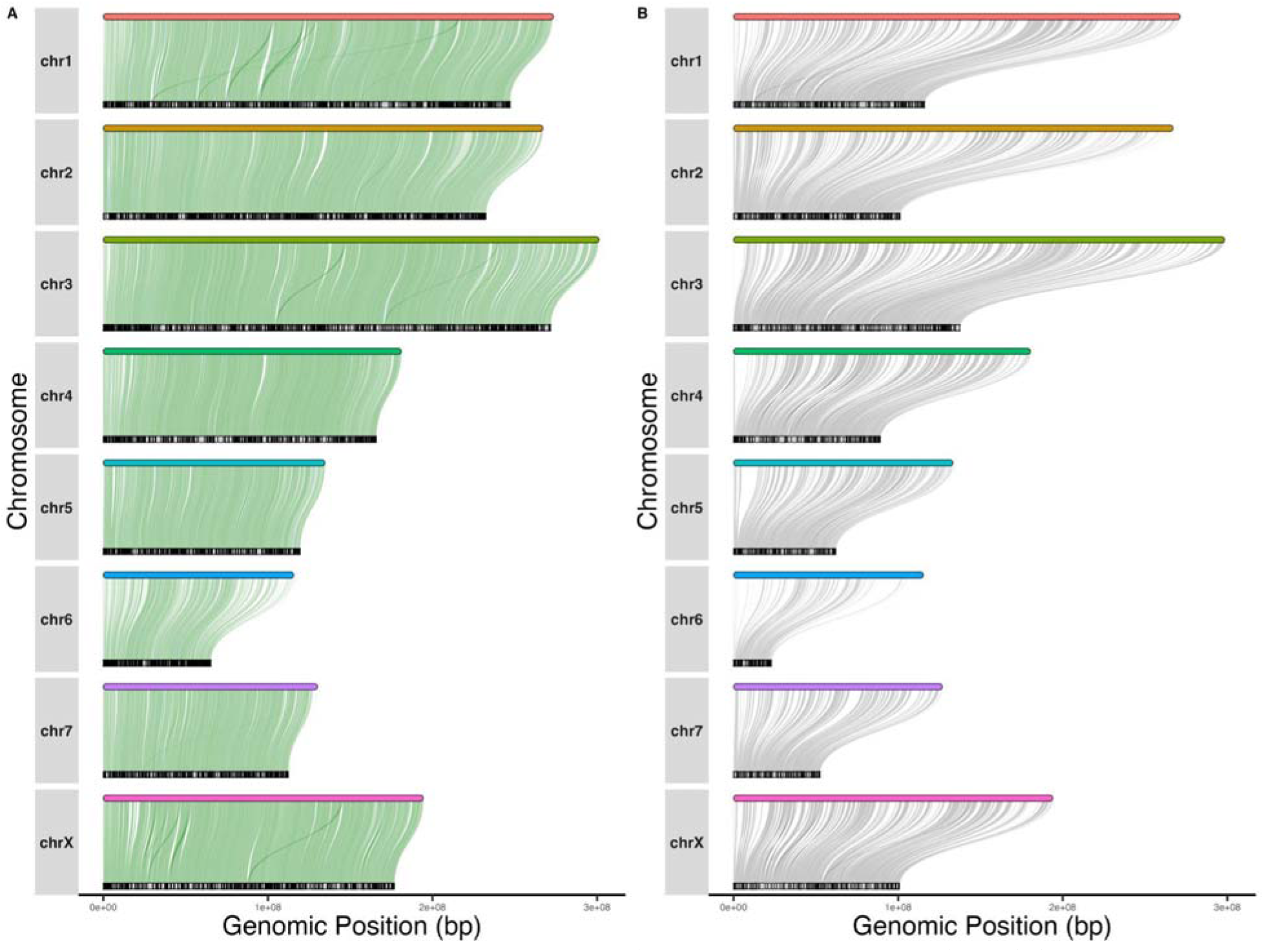
Alignment of all genome scaffolds (top tracks, colored bars) to (A) all scaffolds of the previous genome, as in Figure 1E, or (B) the subset of scaffolds which were placed on pseudochromosomes in the previous genome (bottom tracks, black and grey lines). Note the reduction in the length of each pseudochromosome in (B) compared to (A), indicating the number of unplaced scaffolds in the previous genome. Analysis limited to the scaffolds mapping to the chromosomes from the new assembly and which are 10,000 bp or longer.

**Figure S2.**
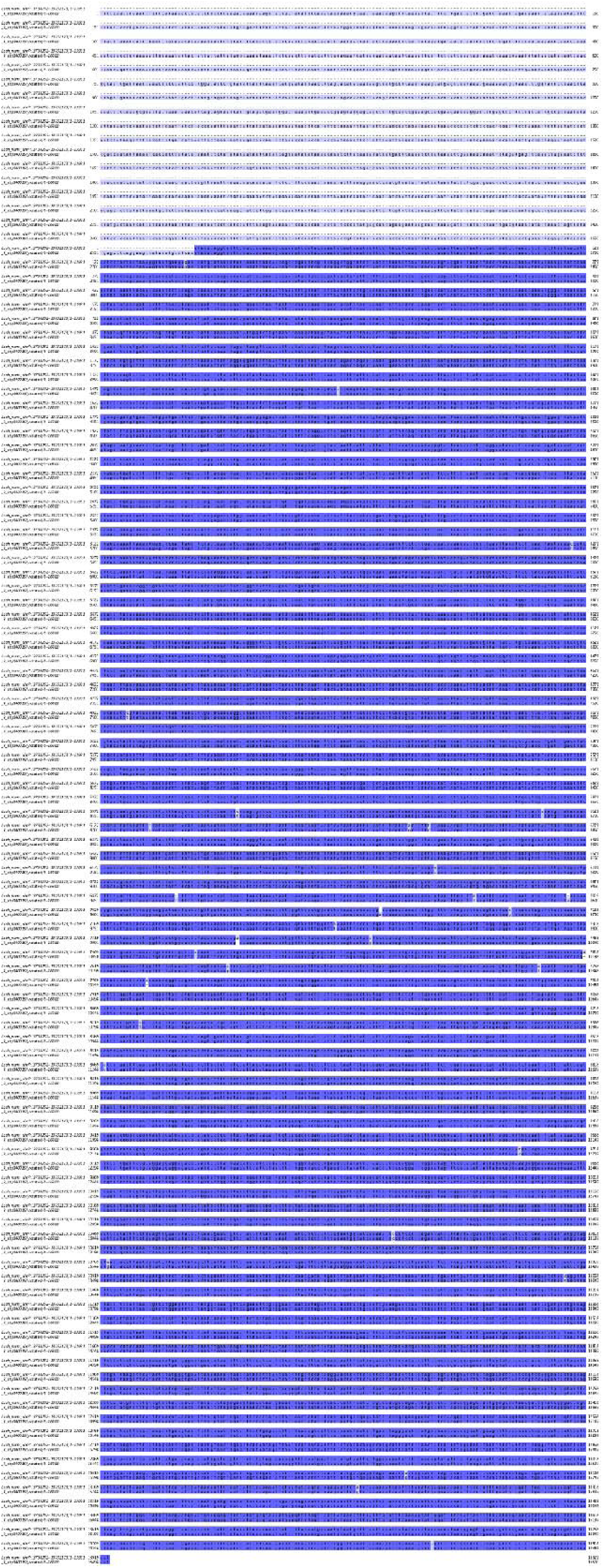
Alignment of the mtDNA against the large NUMT found on chromosome 7. Colored regions are conserved between the two sequences. Sequences have 84.17% similarity, including gaps.

## References

Al Arab, Marwa, Christian Höner Zu Siederdissen, Kifah Tout, Abdullah H. Sahyoun, Peter F. Stadler, and Matthias Bernt. 2017. “Accurate Annotation of Protein-Coding Genes in Mitochondrial Genomes.” Molecular Phylogenetics and Evolution 106 (January): 209–216.

Bao, Weidong, Kenji K. Kojima, and Oleksiy Kohany. 2015. “Repbase Update, a Database of Repetitive Elements in Eukaryotic Genomes.” Mobile DNA 6 (1): 11.

Blankers, Thomas, Kevin P. Oh, Aureliano Bombarely, and Kerry L. Shaw. 2018. “The Genomic Architecture of a Rapid Island Radiation: Recombination Rate Variation, Chromosome Structure, and Genome Assembly of the Hawaiian Cricket Laupala.” Genetics 209 (4): 1329–1344.

Blankers, Thomas, Kevin P. Oh, and Kerry L. Shaw. 2019. “Parallel Genomic Architecture Underlies Repeated Sexual Signal Divergence in Hawaiian Laupala Crickets.” *Proceedings*. Biological Sciences 286 (1912): 20191479.

Blankers, Thomas, and Kerry L. Shaw. 2024. “The Biogeographic and Evolutionary Processes Shaping Population Divergence in Laupala.” Molecular Ecology 33 (15): e17444.

Bowman, Megan. n.d. “Repeat Library Construction-Advanced.” Accessed May 5, 2026. http://weatherby.genetics.utah.edu/MAKER/wiki/index.php/Repeat_Library_Construction-Advanced.

Bukowski, R., Q. Sun, M. Howard, and J. Pillardy. 2010. “BioHPC: Computational Biology Application Suite for High Performance Computing.” Journal of Biomolecular Techniques 21 (3 Suppl): S23–S23.

Camacho, Christiam, George Coulouris, Vahram Avagyan, et al. 2009. “BLAST+: Architecture and Applications.” BMC Bioinformatics 10 (1): 421.

Cantalapiedra, Carlos P., Ana Hernández-Plaza, Ivica Letunic, Peer Bork, and Jaime Huerta-Cepas. 2021. “EggNOG-Mapper v2: Functional Annotation, Orthology Assignments, and Domain Prediction at the Metagenomic Scale.” Molecular Biology and Evolution 38 (12): 5825–5829.

Cechova, Monika. 2020. “Probably Correct: Rescuing Repeats with Short and Long Reads.” Genes 12 (1): 48.

Cheng, Haoyu, Gregory T. Concepcion, Xiaowen Feng, Haowen Zhang, and Heng Li. 2021. “Haplotype-Resolved de Novo Assembly Using Phased Assembly Graphs with Hifiasm.” Nature Methods 18 (2): 170–175.

Chen, Yu, Yixin Zhang, Amy Y. Wang, Min Gao, and Zechen Chong. 2021. “Accurate Long-Read de Novo Assembly Evaluation with Inspector.” Genome Biology 22 (1): 312.

Crescente, Juan Manuel, Diego Zavallo, Marcelo Helguera, and Leonardo Sebastián Vanzetti. 2018. “MITE Tracker: An Accurate Approach to Identify Miniature Inverted-Repeat Transposable Elements in Large Genomes.” BMC Bioinformatics 19 (1): 348.

Cucini, Claudio, Chiara Leo, Nicola Iannotti, et al. 2021. “EZmito: A Simple and Fast Tool for Multiple Mitogenome Analyses.” *Mitochondrial DNA. Part B*, Resources 6 (3): 1101–1109.

Dainat, Jacques, Robrecht Cannoodt, André Soares, et al. 2026. NBISweden/AGAT: AGAT v1.7.0. Zenodo. 10.5281/ZENODO.3552717.

deCarvalho, Tagide N., and Kerry L. Shaw. 2005. “Nuptial Feeding of Spermless Spermatophores in the Hawaiian Swordtail Cricket, Laupala Pacifica (Gryllidae: Triginodiinae).” The Science of Nature 92 (10): 483–487.

Donath, Alexander, Frank Jühling, Marwa Al-Arab, et al. 2019. “Improved Annotation of Protein-Coding Genes Boundaries in Metazoan Mitochondrial Genomes.” Nucleic Acids Research 47 (20): 10543–10552.

Ellinghaus, David, Stefan Kurtz, and Ute Willhoeft. 2008. “LTRharvest, an Efficient and Flexible Software for de Novo Detection of LTR Retrotransposons.” BMC Bioinformatics 9 (1): 18.

Flynn, Jullien M., Robert Hubley, Clément Goubert, et al. 2020. “RepeatModeler2 for Automated Genomic Discovery of Transposable Element Families.” Proceedings of the National Academy of Sciences of the United States of America 117 (17): 9451–9457.

Gabriel, Lars, Tomáš Brůna, Katharina J. Hoff, et al. 2024. “BRAKER3: Fully Automated Genome Annotation Using RNA-Seq and Protein Evidence with GeneMark-ETP, AUGUSTUS, and TSEBRA.” Genome Research 34 (5): 769–777.

Gabriel, Lars, Katharina J. Hoff, Tomáš Brůna, Mark Borodovsky, and Mario Stanke. 2021. “TSEBRA: Transcript Selector for BRAKER.” BMC Bioinformatics 22 (1): 566.

“Galaxy.” n.d. Accessed April 24, 2026. https://usegalaxy.eu/.

Galaxy Community. 2024. “The Galaxy Platform for Accessible, Reproducible, and Collaborative Data Analyses: 2024 Update.” Nucleic Acids Research 52 (W1): W83–W94.

Gillespie, Rosemary G., Gordon M. Bennett, Luc De Meester, et al. 2020. “Comparing Adaptive Radiations across Space, Time, and Taxa.” The Journal of Heredity 111 (1): 1–20.

Grace, Jaime L., and Kerry L. Shaw. 2011. “Coevolution of Male Mating Signal and Female Preference during Early Lineage Divergence of the Hawaiian Cricket, Laupala Cerasina.” Evolution; International Journal of Organic Evolution 65 (8): 2184–2196.

Grace, Jaime L., and Kerry L. Shaw. 2012. “Incipient Sexual Isolation in Laupala Cerasina: Females Discriminate Population-Level Divergence in Acoustic Characters.” Current Zoology 58 (3): 416–425.

Gurevich, Alexey, Vladislav Saveliev, Nikolay Vyahhi, and Glenn Tesler. 2013. “QUAST: Quality Assessment Tool for Genome Assemblies.” *Bioinformatics (Oxford*, England) 29 (8): 1072–1075.

Hawlitschek, Oliver, David Sadílek, Lara-Sophie Dey, et al. 2023. “New Estimates of Genome Size in Orthoptera and Their Evolutionary Implications.” PloS One 18 (3): e0275551.

Hazkani-Covo, Einat, Raymond M. Zeller, and William Martin. 2010. “Molecular Poltergeists: Mitochondrial DNA Copies (numts) in Sequenced Nuclear Genomes.” PLoS Genetics 6 (2): e1000834.

Hiller, Anna E., Michelle S. Koo, Kari R. Goodman, Kerry L. Shaw, Patrick M. O’Grady, and Rosemary G. Gillespie. 2019. “Niche Conservatism Predominates in Adaptive Radiation: Comparing the Diversification of Hawaiian Arthropods Using Ecological Niche Modelling.” Biological Journal of the Linnean Society. Linnean Society of London 127 (2): 479–492.

Huerta-Cepas, Jaime, Damian Szklarczyk, Davide Heller, et al. 2019. “eggNOG 5.0: A Hierarchical, Functionally and Phylogenetically Annotated Orthology Resource Based on 5090 Organisms and 2502 Viruses.” Nucleic Acids Research 47 (D1): D309–D314.

Kataoka, Kosuke, Ryuto Sanno, Tomasz Gaczorek, et al. 2026. “Chromosome-Scale Genome Assembly and Annotation of the Two-Spotted Cricket Gryllus Bimaculatus (Orthoptera: Gryllidae).” G3 (Bethesda, Md.) 16 (4). 10.1093/g3journal/jkag036.

Kataoka, Kosuke, Yuki Togawa, Ryuto Sanno, Toru Asahi, and Kei Yura. 2022. “Dissecting Cricket Genomes for the Advancement of Entomology and Entomophagy.” Biophysical Reviews 14 (1): 75–97.

Krueger, Felix. n.d. TrimGalore: A Wrapper around Cutadapt and FastQC to Consistently Apply Adapter and Quality Trimming to FastQ Files, with Extra Functionality for RRBS Data. Github. Accessed April 20, 2026. https://github.com/felixkrueger/trimgalore.

Kurylo, Cyril, Cervin Guyomar, Sylvain Foissac, and Sarah Djebali. 2023. “TAGADA: A Scalable Pipeline to Improve Genome Annotations with RNA-Seq Data.” NAR Genomics and Bioinformatics 5 (4): lqad089.

Laetsch, Dominik R., and Mark L. Blaxter. 2017. “BlobTools: Interrogation of Genome Assemblies.” F1000Research 6 (July): 1287.

Lang, B. Franz, Natacha Beck, Samuel Prince, Matt Sarrasin, Pierre Rioux, and Gertraud Burger. 2023. “Mitochondrial Genome Annotation with MFannot: A Critical Analysis of Gene Identification and Gene Model Prediction.” Frontiers in Plant Science 14 (July): 1222186.

Li, Heng. 2018. “Minimap2: Pairwise Alignment for Nucleotide Sequences.” *Bioinformatics (Oxford*, England) 34 (18): 3094–3100.

Li, Heng. 2021. “New Strategies to Improve minimap2 Alignment Accuracy.” *Bioinformatics (Oxford*, England) 37 (23): 4572–4574.

Li, Heng. n.d. Seqtk: Toolkit for Processing Sequences in FASTA/Q Formats. Github. Accessed April 17, 2026. https://github.com/lh3/seqtk.

Lima Docs. n.d. “Lima Home.” Accessed April 16, 2026. https://lima.how/.

Liu, Qian, Peng Zhang, Depeng Wang, Weihong Gu, and Kai Wang. 2017. “Interrogating the ‘Unsequenceable’ Genomic Trinucleotide Repeat Disorders by Long-Read Sequencing.” Genome Medicine 9 (1): 65.

Liu, Xuanzeng, Nian Liu, Xuan Jing, et al. 2024. “Genomic and Transcriptomic Perspectives on the Origin and Evolution of NUMTs in Orthoptera.” Molecular Phylogenetics and Evolution 201 (108221): 108221.

Liu, Xuanzeng, Lina Zhao, Muhammad Majid, and Yuan Huang. 2024. “Orthoptera-TElib: A Library of Orthoptera Transposable Elements for TE Annotation.” Mobile DNA 15 (1): 5.

Manni, Mosè, Matthew R. Berkeley, Mathieu Seppey, Felipe A. Simão, and Evgeny M. Zdobnov. 2021. “BUSCO Update: Novel and Streamlined Workflows along with Broader and Deeper Phylogenetic Coverage for Scoring of Eukaryotic, Prokaryotic, and Viral Genomes.” Molecular Biology and Evolution 38 (10): 4647–4654.

Meier, Joana I., Matthew D. McGee, David A. Marques, et al. 2023. “Cycles of Fusion and Fission Enabled Rapid Parallel Adaptive Radiations in African Cichlids.” *Science (New York*, N.Y.) 381 (6665): eade2833.

Mendelson, Tamra C., and Kerry L. Shaw. 2005. “Sexual Behaviour: Rapid Speciation in an Arthropod.” Nature 433 (7024): 375–376.

Mullen, Sean P., Tamra C. Mendelson, Coby Schal, and Kerry L. Shaw. 2007. “Rapid Evolution of Cuticular Hydrocarbons in a Species Radiation of Acoustically Diverse Hawaiian Crickets (Gryllidae: Trigonidiinae: Laupala).” Evolution; International Journal of Organic Evolution 61 (1): 223–231.

Oh, Kevin P., and Kerry L. Shaw. 2013. “Multivariate Sexual Selection in a Rapidly Evolving Speciation Phenotype.” *Proceedings*. Biological Sciences 280 (1761): 20130482.

Otte, Daniel. 1994. *The Crickets of Hawaii*. Listening Chamber Poetry Series. Orthopterists’ Society.

Paysan-Lafosse, Typhaine, Matthias Blum, Sara Chuguransky, et al. 2023. “InterPro in 2022.” Nucleic Acids Research 51 (D1): D418–D427.

Peart, Claire R., Christina Williams, Saurabh D. Pophaly, et al. 2021. “Hi-C Scaffolded Short- and Long-Read Genome Assemblies of the California Sea Lion Are Broadly Consistent for Syntenic Inference across 45 Million Years of Evolution.” Molecular Ecology Resources 21 (7): 2455–2470.

Pedersen, Brent S., and Aaron R. Quinlan. 2018. “Mosdepth: Quick Coverage Calculation for Genomes and Exomes.” *Bioinformatics (Oxford*, England) 34 (5): 867–868.

Petrov, D. A., T. A. Sangster, J. S. Johnston, D. L. Hartl, and K. L. Shaw. 2000. “Evidence for DNA Loss as a Determinant of Genome Size.” *Science (New York*, N.Y.) 287 (5455): 1060– 1062.

Quinlan, Aaron R. 2014. “BEDTools: The Swiss-Army Tool for Genome Feature Analysis.” Current Protocols in Bioinformatics 47 (1): 11.12.1–34.

Rundell, Rebecca J., and Trevor D. Price. 2009. “Adaptive Radiation, Nonadaptive Radiation, Ecological Speciation and Nonecological Speciation.” Trends in Ecology & Evolution 24 (7): 394–399.

Schluter, Dolph. 2000. The Ecology of Adaptive Radiation. Oxford Series in Ecology and Evolution. Oxford University Press.

Scholl, Joshua P., and John J. Wiens. 2016. “Diversification Rates and Species Richness across the Tree of Life.” *Proceedings*. Biological Sciences 283 (1838): 20161334.

Sen, Raunak, Nicholai M. Hensley, Bhaavya Srivastava, Wout van der Heide, and Kerry L. Shaw. 2025. “Sexual Isolation Maintains Species Boundaries between Hawaiian Crickets in Sympatry despite Weak Habitat Isolation.” In *bioRxiv*. BioRxiv, September 8. 10.1101/2025.09.04.674279.

Sen, Raunak, and Kerry L. Shaw. 2025. “Genomic Data Confirms Phenotypic Predictions of Hybridization between Cryptic Hawaiian Cricket Species.” In bioRxiv. September 15. 10.1101/2025.09.09.675270.

Shaw, Kerry L. 2000. “Further Acoustic Diversity in Hawaiian Forests: Two New Species of Hawaiian Cricket (Orthoptera: Gryllidae: Trigonidiinae: Laupala).” Zoological Journal of the Linnean Society 129 (1): 73–91.

Shaw, Kerry L. 2002. “Conflict between Nuclear and Mitochondrial DNA Phylogenies of a Recent Species Radiation: What mtDNA Reveals and Conceals about Modes of Speciation in Hawaiian Crickets.” Proceedings of the National Academy of Sciences of the United States of America 99 (25): 16122–16127.

Shaw, Kerry L., and Albert H. Khine. 2004. “Courtship Behavior in the Hawaiian Cricket *Laupala Cerasina*: Males Provide Spermless Spermatophores as Nuptial Gifts.” Ethology: Formerly Zeitschrift Für Tierpsychologie 110 (2): 81–95.

Shaw, Kerry L., and Sky C. Lesnick. 2009. “Genomic Linkage of Male Song and Female Acoustic Preference QTL Underlying a Rapid Species Radiation.” Proceedings of the National Academy of Sciences of the United States of America 106 (24): 9737–9742.

Shaw, Kerry L., Yvonne M. Parsons, and Sky C. Lesnick. 2007. “QTL Analysis of a Rapidly Evolving Speciation Phenotype in the Hawaiian Cricket Laupala.” Molecular Ecology 16 (14): 2879–2892.

Shin, Seunggwan, Austin J. Baker, Jacob Enk, et al. 2024. “Orthoptera-Specific Target Enrichment (OR-TE) Probes Resolve Relationships over Broad Phylogenetic Scales.” Scientific Reports 14 (1): 21377.

Shumate, Alaina, and Steven L. Salzberg. 2019. “Liftoff: A Pipeline for Mapping Genes onto High-Quality Genome Assemblies.” November 11. 10.7490/f1000research.1117639.1.

Skera Docs. n.d. “Skera Home.” Accessed April 16, 2026. https://skera.how/.

Song, Hojun, Olivier Béthoux, Seunggwan Shin, et al. 2020. “Phylogenomic Analysis Sheds Light on the Evolutionary Pathways towards Acoustic Communication in Orthoptera.” Nature Communications 11 (1): 4939.

Song, Hojun, Matthew J. Moulton, and Michael F. Whiting. 2014. “Rampant Nuclear Insertion of mtDNA across Diverse Lineages within Orthoptera (Insecta).” PloS One 9 (10): e110508.

Soria-Carrasco, Víctor, Zachariah Gompert, Aaron A. Comeault, et al. 2014. “Stick Insect Genomes Reveal Natural Selection’s Role in Parallel Speciation.” *Science (New York*, N.Y.) 344 (6185): 738–742.

Steinbiss, Sascha, Ute Willhoeft, Gordon Gremme, and Stefan Kurtz. 2009. “Fine-Grained Annotation and Classification of de Novo Predicted LTR Retrotransposons.” Nucleic Acids Research 37 (21): 7002–7013.

Storer, Jessica, Robert Hubley, Jeb Rosen, Travis J. Wheeler, and Arian F. Smit. 2021. “The Dfam Community Resource of Transposable Element Families, Sequence Models, and Genome Annotations.” Mobile DNA 12 (1): 2.

Szrajer, Szymon, David Gray, and Guillem Ylla. 2024. “The Genome Assembly and Annotation of the Cricket Gryllus Longicercus.” Scientific Data 11 (1): 708.

TransDecoder: TransDecoder Source. n.d. Github. Accessed April 19, 2026. https://github.com/TransDecoder/TransDecoder.

“TransposonPSI: An Application of PSI-Blast to Mine (Retro-)Transposon ORF Homologies.” n.d. Accessed April 17, 2026. https://transposonpsi.sourceforge.net/.

Uliano-Silva, Marcela, João Gabriel R. N. Ferreira, Ksenia Krasheninnikova, et al. 2023. “MitoHiFi: A Python Pipeline for Mitochondrial Genome Assembly from PacBio High Fidelity Reads.” BMC Bioinformatics 24 (1): 288.

UniProt Consortium. 2023. “UniProt: The Universal Protein Knowledgebase in 2023.” Nucleic Acids Research 51 (D1): D523–D531.

University of Greifswald. 2018. “Bioinformatics Web Server - University of Greifswald.” July 17. https://bioinf.uni-greifswald.de/bioinf/partitioned_odb11/.

Vassetzky, Nikita S., and Dmitri A. Kramerov. 2013. “SINEBase: A Database and Tool for SINE Analysis.” Nucleic Acids Research 41 (Database issue): D83–9.

Vurture, Gregory W., Fritz J. Sedlazeck, Maria Nattestad, et al. 2017. “GenomeScope: Fast Reference-Free Genome Profiling from Short Reads.” *Bioinformatics (Oxford*, England) 33 (14): 2202–2204.

Waller, Hayden, Thomas Blankers, Mingzi Xu, and Kerry L. Shaw. 2023. “Quantitative Trait Loci Underlying a Speciation Phenotype.” Insect Molecular Biology 32 (6): 592–602.

Wei, Wei, Katherine R. Schon, Greg Elgar, et al. 2022. “Nuclear-Embedded Mitochondrial DNA Sequences in 66,083 Human Genomes.” Nature 611 (7934): 105–114.

Wen, Huaming, Jinbao Yang, Xianjia Zhao, et al. 2025. “TRFill: Synergistic Use of HiFi and Hi-C Sequencing Enables Accurate Assembly of Tandem Repeats for Population-Level Analysis.” Genome Biology 26 (1): 227.

Wiens, John J., and Daniel S. Moen. 2025. “Rapid Radiations Underlie Most of the Known Diversity of Life.” Frontiers in Ecology and Evolution 13 (1596591). 10.3389/fevo.2025.1596591.

Wiley, Chris, Christopher K. Ellison, and Kerry L. Shaw. 2012. “Widespread Genetic Linkage of Mating Signals and Preferences in the Hawaiian Cricket Laupala.” *Proceedings*. Biological Sciences 279 (1731): 1203–1209.

Xu, Mingzi, and Kerry L. Shaw. 2019. “Genetic Coupling of Signal and Preference Facilitates Sexual Isolation during Rapid Speciation.” *Proceedings*. Biological Sciences 286 (1913): 20191607.

Xu, Mingzi, and Kerry L. Shaw. 2020. “Spatial Mixing between Calling Males of Two Closely Related, Sympatric Crickets Suggests Beneficial Heterospecific Interactions in a NonAdaptive Radiation.” The Journal of Heredity 111 (1): 84–91.

Xu, Mingzi, and Kerry L. Shaw. 2021. “Extensive Linkage and Genetic Coupling of Song and Preference Loci Underlying Rapid Speciation in Laupala Crickets.” The Journal of Heredity 112 (2): 204–213.

Xu, Mingzi, and Kerry L. Shaw. 2026. “Linked Song and Preference Loci Suggests Substantial Contribution of Genetic Coupling in Rapid Speciation of the Laupala Crickets.” Genetics, no. iyag073 (March). 10.1093/genetics/iyag073.

Ylla, Guillem, Taro Nakamura, Takehiko Itoh, et al. 2021. “Insights into the Genomic Evolution of Insects from Cricket Genomes.” Communications Biology 4 (1): 733.

Zhang, Shangzhe, Kristin R. Duffield, Bert Foquet, et al. 2025. “A High-Quality Reference Genome and Comparative Genomics of the Widely Farmed Banded Cricket (Gryllodes Sigillatus) Identifies Selective Breeding Targets.” Ecology and Evolution 15 (3): e71134.

Zhou, Weichen, Kalpita R. Karan, Wenjin Gu, et al. 2024. “Somatic Nuclear Mitochondrial DNA Insertions Are Prevalent in the Human Brain and Accumulate over Time in Fibroblasts.” PLoS Biology 22 (8): e3002723.

